# Engineering bacterial blight-resistant plants through CRISPR/Cas9-targeted editing of the *MeSWEET10a* promoter in cassava

**DOI:** 10.1101/2022.03.02.482644

**Authors:** Yajie Wang, Mengting Geng, Ranran Pan, Tong Zhang, Xiaohua Lu, Xinghou Zhen, Yannian Che, Ruimei Li, Jiao Liu, Yinhua Chen, Jianchun Guo, Yuan Yao

## Abstract

Cassava starch is a widely used raw materials for industrial production. However, cassava bacterial blight (CBB) caused by *Xanthomonas axonopodis* pv. *manihotis* (*Xam*) results in severe yield losses and is the most destructive bacterial disease in all worldwide cassava planting regions. This study showed that editing of the promoter in the disease-susceptibility gene *MeSWEET10a* of SC8 cassava confers resistance to CBB. All mutated cassava lines had normal morphological and yield-related traits as the wild type. The results lay a research foundation for breeding cassava resistant to bacterial blight.

**Highlights:** *MeSWEET10a* gene in SC8 cultivar is hijacked by TALE20 from *Xam11* strain.

Editing of the *MeSWEET10a* promoter in SC8 cultivar confers resistance to CBB.

All the mutated cassava lines had similar yield-related traits compared to wild-type.

## 1. Introduction

Starch is one of the most important polymers that has been extensively used in industry. Cassava is a major available source of commercially produced starches in the market. It is extensively cultivated throughout the tropical and subtropical regions. However, cassava bacterial blight (CBB) caused by *Xanthomonas axonopodis* pv. *manihotis* (*Xam*) results in severe yield losses and is the most destructive bacterial disease in all worldwide cassava planting regions (McCallum et al., 2017). The vast majority of cassava cultivars are not resistant to CBB. *Xam* pathogen infection secretes transcription activator-like effectors (TALEs) into cassava host cells through the type III secretion system (Boch and Bonas, 2010). The repeat variable diresidues (RVDs) in TALEs recognize the effector-binding elements (EBEs) on the host susceptibility (S) gene promoters and activate S gene expression which facilitates bacterial growth and disease symptom formation. *Xam668* is a highly virulent *Xam* strain to cassava which contains five transcription activator-like effectors (TALEs) including TALE20_*Xam668*_ (Bart et al., 2012). The repeat variable diresidues (RVDs) in TALE20_*Xam668*_ recognize the effector-binding element (EBE) on the *MeSWEET10a* promoter from cassava 60444 cultivar and activate *MeSWEET10a* gene expression which facilitates sucrose transport from the interior of plant cells to the apoplasts and provide carbon sources for the bacteria (Cohn et al., 2014). Thus, editing the EBE in the *MeSWEET10a* promoter may potentially confer resistance to CBB in cassava cultivars.

*Xam11* is a highly pathogenic subspecies from Hainan, China that infects the Chinese local planting cassava SC8 cultivar (*Manihot esculenta* Crantz South China 8) with marked pathogenic symptoms. In this study, the CRISPR/Cas9 system was used to edit the EBE_TALE20_ of the *MeSWEET10a* promoter in SC8 cultivar. The EBE_TALE20_ mutant plants showed enhanced resistance to *Xam11*, and did not show any significant differences in major yield-related traits compared to wild-type plants.

## 2. Materials and Methods

### 2.1. Plant Materials

For cassava transformation, the SC8 cultivar was used as the recipient. For pathogen inoculation, SC8 and CRISPR/Cas9 mutant plants were cultivated in a greenhouse under a 16-h/8-h light/dark cycle at 23-28□. For agronomic trait (weight and number of tuber root per plant, dry matter ratio, starch content) evaluation, all of the tested mutant plants and WT were planted in the 15-cm round pots containing 10% Perlite, 80% soil.

### 2.2. Dual LUC Reporter Gene Assay

Genomic DNA extraction from SC8 leaves was performed according to the Plant DNA Kit. The promoter sequence of *MeSWEET10a* from SC8 was amplified by using the specific primers (F: 5’-AACTTTTAGAATGAGCCCTTG-3’; R: 5’-TTCTCCGGCTATAGTAGAGAC-3’), then inserted into pGreenII0800-LUC vector, named as pGreen II0800-pMeSWEET10a-LUC vector. The EBE_TALE20_ from the *MeSWEET10a* promoter was deleted and generated pSWEET10a-EBE, then inserted into pGreenII0800-LUC, named as pGreenII0800-pMeSWEET10a-EBE_TALE20_-LUC vector. Full-length CDS of TALE20_*xam11*_ was inserted into pGreen II62-SK vector to generate pGreen II62-SK-TALE20_*Xam11*_ vector. The vectors of pGreen II62-SK or pGreen II62-SK-TALE20_*Xam11*_ were used as the effectors, and pGreen II0800-pMeSWEET10a-LUC or pGreen II0800-pMeSWEET10a-EBE_TALE20_-LUC were used as the reporters. The effector and reporter vectors were transformed into tobacco leaves as previously described (Liu et al., 2019b). The activity of Firefly luciferase and the reference Renilla luciferase were quantified using the Dual-Luciferase Reporter Assay System (Promega), according to the manufacturer’s instructions. Three biological repeats were analyzed.

### 2.3. Editing of MeSWEET10a EBE_TALE20_ by CRISPR/Cas9 System

In our previous study, the target sgRNA of *MeSWEET10a* EBE_TALE20_ has been designed and constructed the vector of pCAMBIA1301-Cas9-EBE-sgRNA, and the editing function of target sgRNA has been validated in cassava FEC by transient transformation (Zhang et al., 2021). In our current research, the friable embryogenic callus (FEC) of SC8 cassava was infected by *Agrobacterium* LBA4404 harboring editing vector of pCAMBIA1301-Cas9-EBE-sgRNA. The transgenic positive lines were produced according to the protocol described by Nyaboga et al. (2015) (Nyaboga et al., 2015). The transformation conditions of cassava SC8 were *Agrobacterium* strain LBA4404 cell infection (density OD_600_ = 0.65), 250 μM acetosyringone induction, and agro-cultivation with wet FEC for 3 days in dark. To analyze the genotypes of the CRISPR/Cas9 mutants, genomic DNA from the transgenic positive lines were extracted, and analyzed using the Hi-TOM program for high-throughput mutations (Liu et al., 2019a).

### 2.4. Plant Inoculation Xam11 and Symptom Observation

*Xanthomonas* strain *Xam11* was resuspended in 10 mM MgCl_2_. For watersoaking assays and qRT-PCR analyses, the leaves of SC8 and mutant lines (#1, #2, #3, #4) were injected by *Xam11* via a 1-mL needleless syringe. 5 μL of the bacterial suspension at OD_600_ = 0.1 density was injected to the back side of leaves. Three biological repeats were analyzed. The expression of *MeSWEET10a* gene was measured at 0, 0.5, 1, 3, and 7 dpi time points. Symptom development after *Xam11* inoculation was photographed at 3, 7, 14, and 40 dpi. The leaf disease areas after *Xam11* inoculation were measured at 3, 7, and 14 dpi. Bacterial numbers of *Xam11* in the SC8 and mutant lines were counted according to the method of Li et al. (2018)(Li et al., 2018).

### 2.5. qRT-PCR

To analyze *MeSWEET10a* expressions in SC8 cassava and mutant lines, total RNAs isolated from the leaves of SC8 and mutant lines (#1, #2, #3, #4) using RNAplant Plus reagent (TianGen, Beijing, China) following the manufacturer’s instructions were reversed by using the reverse transcriptase kit (TaKaRa, Dalian, China). The qRT-PCR was performed with the reversed cDNAs as substrates and the *MeSWEET10a* specific primers (F: 5’-TCCTCACCTTGACTGCGCTG-3’; R: 5’-AGCACCATCTGGACAATCCCA-3’) by using the SYBR Premix Taq Kit (TaKaRa, Dalian, China) in the ABI7500 Real-Time PCR System (Applied Biosystems, USA). The expression of cassava tubulin gene (Phytozome name: Manes.08G061700) was used as an internal standard by using the primers (F: 5’-GTGGAGGAACTGGTTCTGGA-3’; R: 5’-TGCACTCATCTGCATTCTCC-3’). The 2^-*ΔΔ*Ct^ method was used for relative quantification (Livak and Schmittgen, 2001). Expression data were collected from three biological repeats.

### 2.6. Statistical Analyses

All the data were presented as mean ± *SD* from at least three independent experiments with three replicates. Statistical analyses were conducted by using Student’ s t-tests.

## 3. Results and discussion

### 3.1. MeSWEET10a Gene in SC8 Cultivar is Hijacked by TALE20_Xam11_ from Xam11 Strain

*TALE20* was cloned based on the genome sequencing information of *Xam11*. TALE20_*Xam11*_ (TALE20 in *Xam11*) has 20 RVD with variable amino acids at positions 12 and 13 in each repeat, and EBE_TALE20_ is targeted by TALE20_*Xam11*_ and is located in the −92 to −112 bp region before the *MeSWEET10a* start codon (Fig. 1A). The dual luciferase reporter assay demonstrated that TALE20_*Xam11*_ activated the *MeSWEET10a* promoter from SC8 cultivar and initiated downstream expression of firefly luciferase; however, TALE20_*Xam11*_ could not activate the *MeSWEET10a* promoter lacking the EBE_TALE20_ region (Fig. 1B). This result suggests that the transcription activator-like effector TALE20_*Xam11*_ of *Xam11* strain regulates the expression of *MeSWEET10a* genes by binding to the EBE_TALE20_ region of SC8 cultivar.

**Fig. 1.**
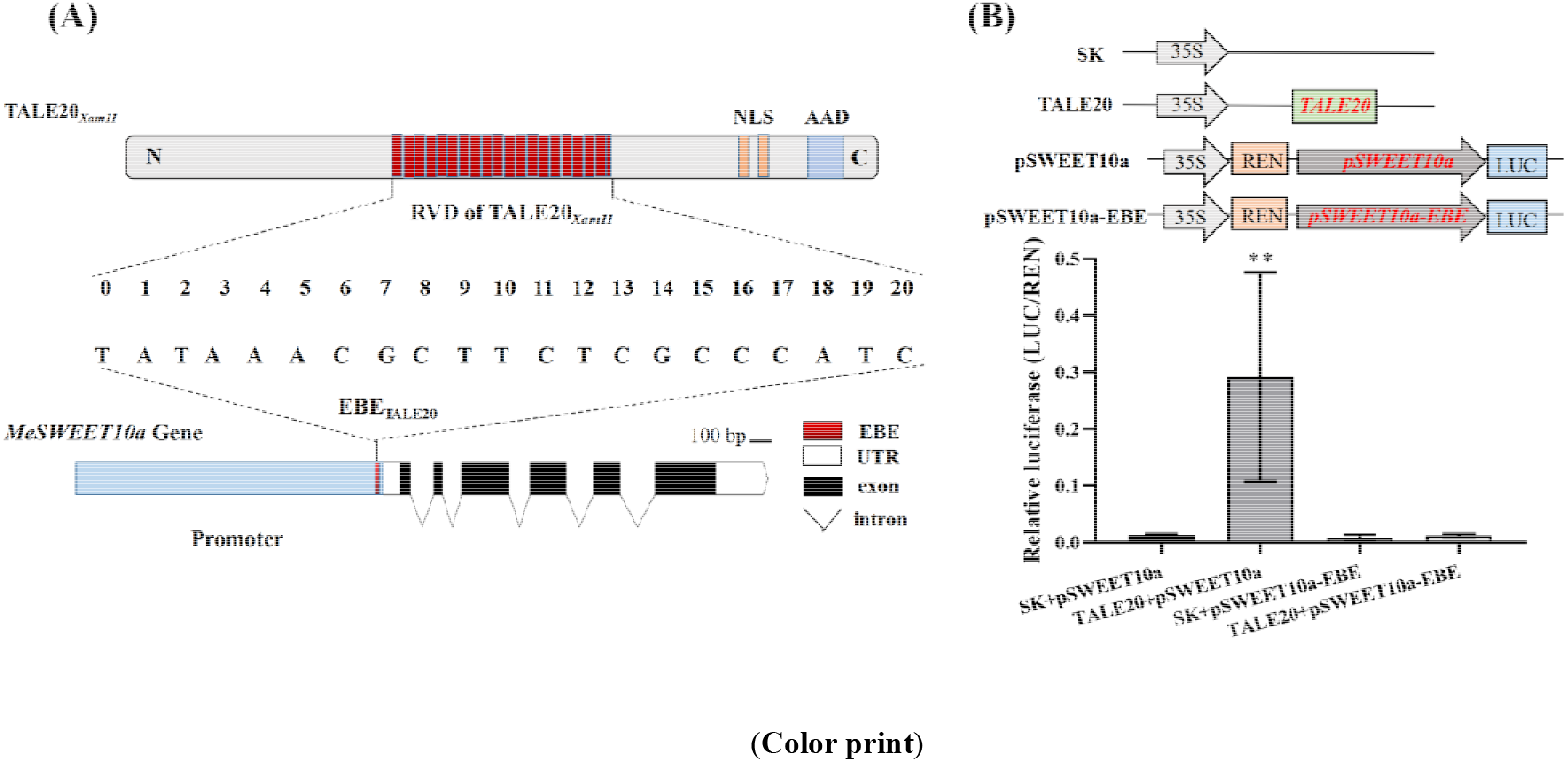
Schematic representation of RVD in TALE20_*Xam11*_ recognize the EBE_*Xam11*_ on the *MeSWEET10a* promoter from cultivar cassava SC8 **(A)**. The C-terminus of TALE20_*Xam11*_ has two nuclear localization signals (NLS) and an acidic activation domain (ADD). TALE20_*Xam11*_ directly activates *MeSWEET10a* expression through the EBE region **(B)**. SK: pGreen II62-SK vector (control vector); TALE20: pGreen II62-SK-TALE20_*Xam11*_ vector; pSWEET10a: pGreen II0800-pMeSWEET10a-LUC vector; pSWEET10a-EBE: pGreen II0800-pMeSWEET10a-EBE_TALE20_-LUC vector (deletion of EBE_TALE20_ from the *MeSWEET10a* promoter). Data are displayed from three biological repeats (Student’s *t*-test, \*\**P* < 0.01).

### 3.2. CRISPR/Cas9-Mediated Editing of EBE_TALE20_ in the Promoter of MeSWEET10a

Inhibition of S gene expression by genetic engineering is an attractive strategy to improve plant disease resistance (Zaidi et al., 2018). Disruption of EBEs in promoters of three major target rice *SWEET* genes by CRISPR-Cas9 avoids recognition by TALEs from *Xanthomonas oryzae* pv. *oryzae* (*Xoo*) and improves bacterial blight resistance (Xu et al., 2019). In this study, the effect of genetically modifying the EBE in the *MeSWEET10a* promoter on CBB resistance after *Xam11* infection was investigated in SC8 cassava. The pCAMBIA1301-Cas9-EBE-sgRNA plasmid was constructed to target EBE_TALE20_ of the *MeSWEET10a* promoter and transformed into the SC8 cassava genome by *Agrobacterium*-mediated fragile embryonic callus (EFC) transformation. In total, 57 independent transgenic lines were regenerated. The EBE_TALE20_ regions from all the transgenic lines were subjected to high throughput tracking of mutation (Hi-TOM) sequences, the results showed that Cas9 activity efficiency was 98.2%. Three types of mutations were obtained among the transgenic lines: homozygous mono-allelic (19.30%), homozygous bi-allelic (61.40%), and heterozygous (17.54%) (Fig. 2A). The EBE_TALE20_ regions in the four mutant types of transgenic cassava lines (#1, #2, #3, #4) and wild type SC8 cassava (WT) were shown in Fig. 2B. These studies showed that the gene editing plasmid could efficiently edit the EBE region of *MeSWEET10a* promoter in SC8 cassava and obtained high proportion of homozygous mutant lines. Up to now, there were five genes (*nCBP-1*, *nCBP-2*, *GBSS*, *PTST1*, and *MePDS*) have been mutated in cassava by CRISPR/Cas9 technology; however, the homozygous mutations in these genes were very low (Odipio et al., 2017; Bull et al., 2018; Gomez et al., 2019).

**Fig. 2.**
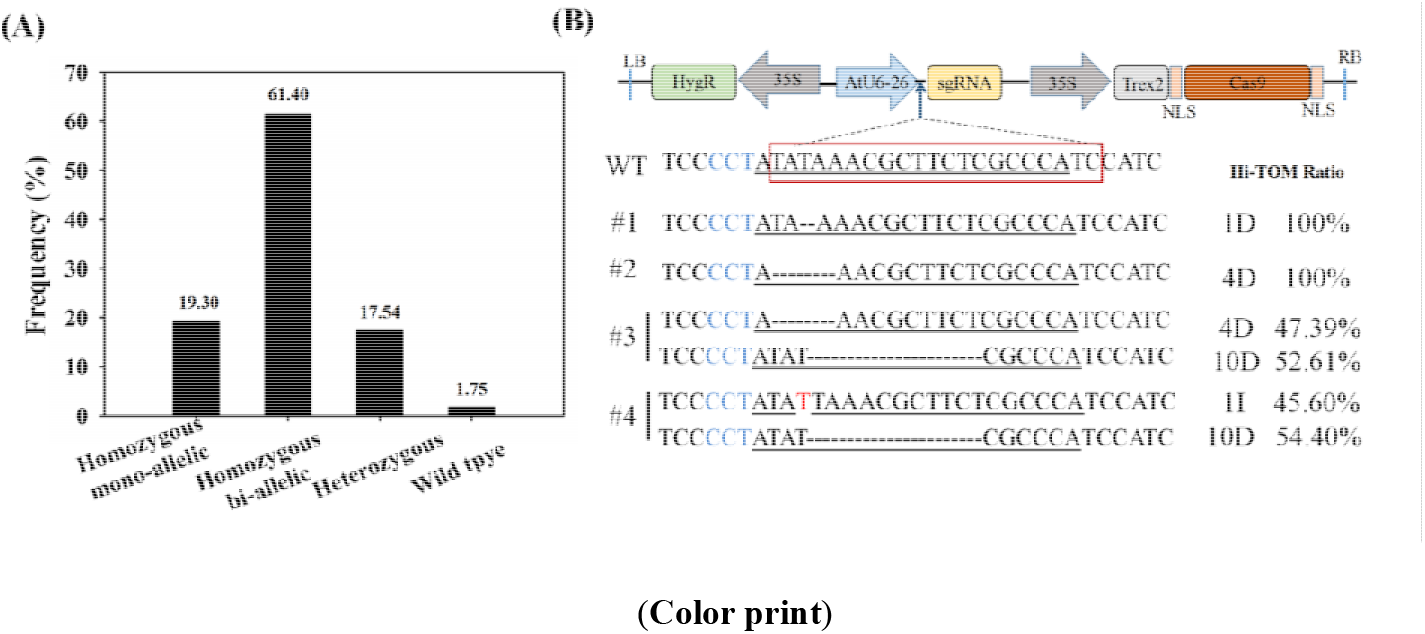
CRISPR/Cas9-induced mutation types and frequency **(A)**. Hi-TOM sequencing of the EBE_TALE20_ regions of wild type (WT) non-transgenic SC8 cassava compared with the engineered lines #1, #2, #3, and #4 **(B)**. The EBE_TALE20_ region of the TALE20_*Xam11*_ target is marked in a red box in WT; the sgRNA sequences are underlined; the protospacer adjacent motif (PAM) sites are marked in blue text; the hyphens are missing bases, and the red letter is the inserted base.

### 3.3. The EBE_TALE20_ Mutant Shows Enhanced Resistance to CBB

The effects of EBE_TALE20_ modification on *MeSWEET10a* expression in the mutant cassava lines were further investigated after inoculation with *Xam11*. The relative expression levels of *MeSWEET10a* in the mutant lines showed no obvious difference from that of the wild type at 0 and 0.5 day post-inoculation (dpi); however, markedly lower levels were observed in the mutant lines compared with that of the wild type at 1–7 dpi (Fig. 3), which showed that EBE_TALE20_ mutations repressed *MeSWEET10a* expression after *Xam11* infection. The wild type had obvious water-soaked symptoms at 3 dpi, however, which were not observed in the mutant lines (#1, #2, #3, #4). At 7 dpi, the wild type had obvious pustule symptoms, while the mutant lines showed slightly water-soaked symptoms; at 14 dpi, the injection sites in the wild type were withered and turned yellow, while in the mutant lines, the injection sites started to wither and the plaque expanded slowly; at 40 dpi, the whole leaves in the wild type were withered and become yellow, whereas the leaves in the mutant lines only withered near the plaques (Fig. 4A). Diseased lesions in the mutant lines were significantly smaller at 3 dpi, 7 dpi, and 14 dpi than that in the wild type (Fig. 4B), it suggested that EBE_TALE20_ mutations limited the expansion of CBB symptoms in cassava SC8. The bacterial growth assay showed that *Xam11* populations in the four mutant lines were significantly lesser than that in the wild type at 3 dpi and 7 dpi (Fig. 4C). These results revealed that EBE_TALE20_ modification of *MeSWEET10a* in the four edited cassava lines had strong and stable resistance to CBB.

**Fig. 3.**
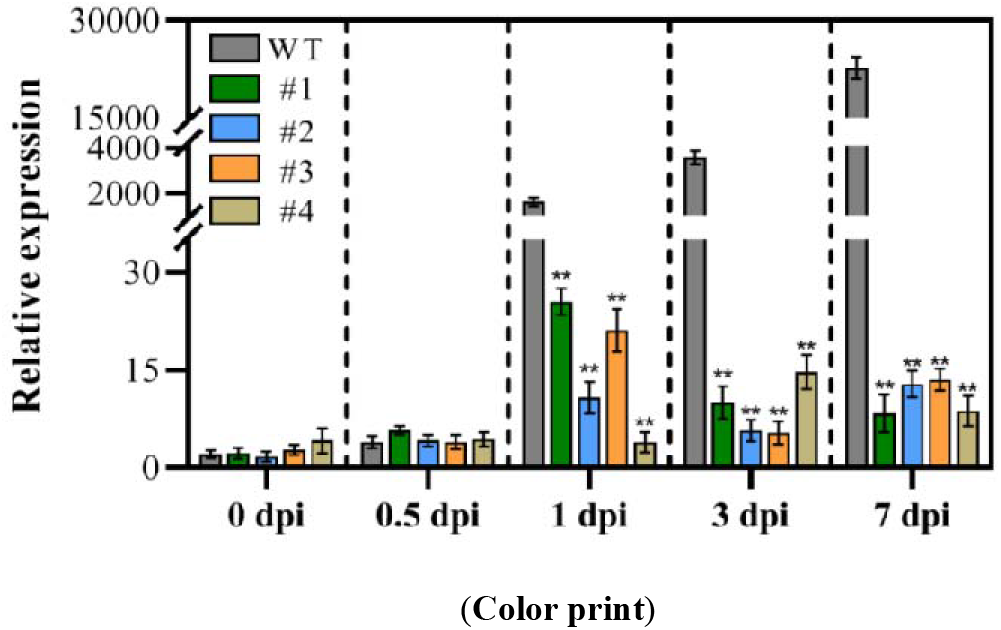
Time course of *MeSWEET10a* expression in mutant lines and WT after *Xam11* inoculation. Three biological repeats were performed for each data point. Statistical tests were two-sided using Student’s t-test compared to WT (\*\**p* < 0.01).

**Fig. 4.**
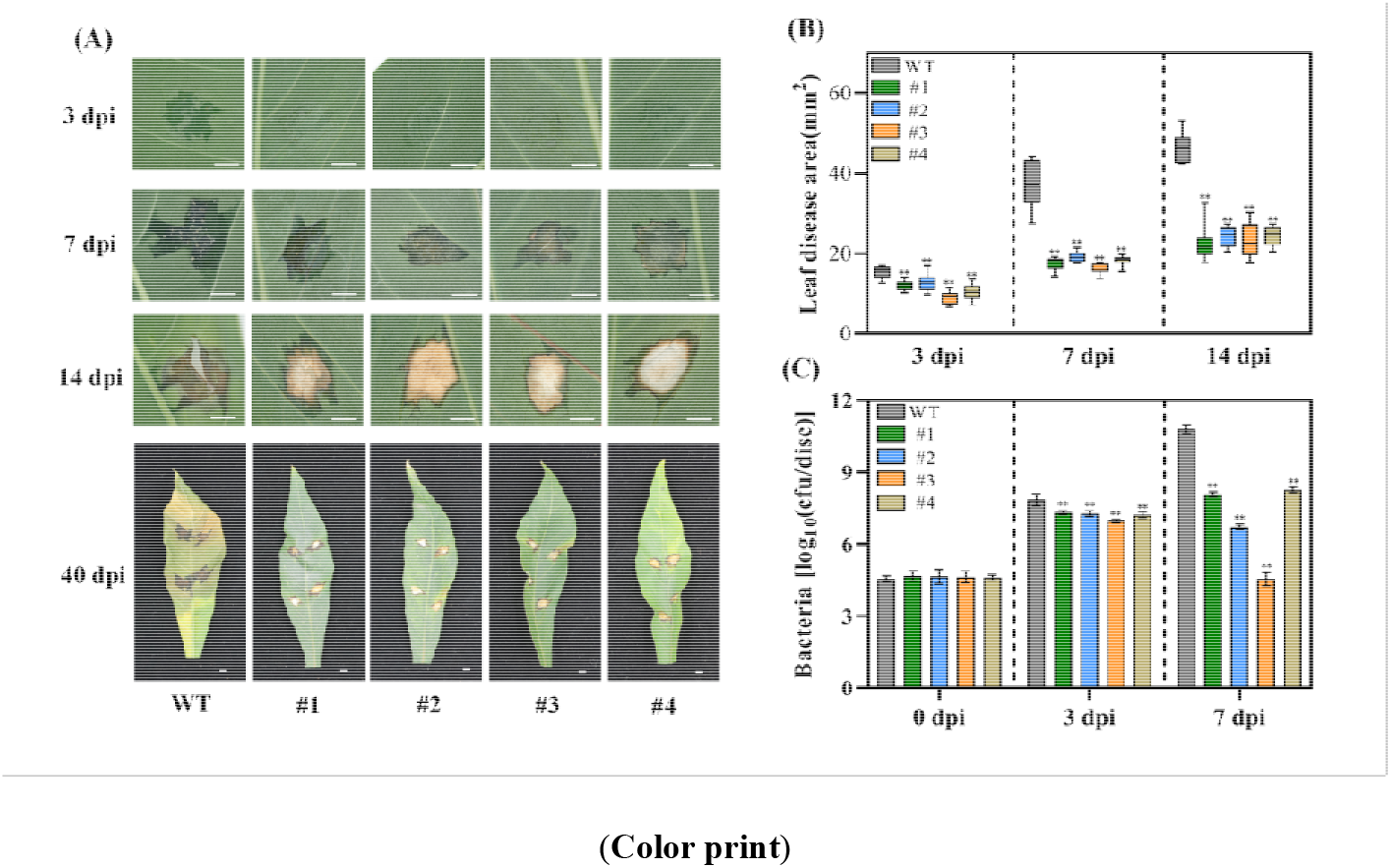
Symptom development after *Xam11* inoculation **(A)**. Scale bar, 5 mm. Time course of leaf disease areas **(B)** and bacterial numbers **(C)** after *Xam11* inoculation. At least 10 leaves were harvested for the diseased areas with the bacterial number assay performed using three biological replicates. Statistical tests were two-sided using Student’s *t*-test compared to WT (\*\**p* < 0.01).

### 3.4. The EBE_TALE20_ Mutant Shows Yield-related Traits Similar to the Wild Type

Mutation of *Mildew resistance locus O* (*Mlo*) through CRISPR-Cas9 has conferred PM resistance in wheat (Wang et al., 2014). In addition, mutations of other S genes such as *eIF4E* and *MPK* enhances disease resistance (Gomez et al., 2019; Xie and Yang, 2013). However, these mutations often affect plant fitness with reduced growth, fertility, yield, and abiotic stress tolerance (Zaidi et al., 2018). Recently, Veley et al. (2022) has been reported that directed methylation to EBE_TALE20_ region of *MeSWEET10a* promoter of TME419 in 60444 cassava cultivar, the plants displayed an increased resistance to *Xam668* strain, meanwhile, they maintained normal growth and development (Veley et al., 2022). In our results, there were no significant differences in tuber root traits and starch content between the EBE_TALE20_ mutated lines and the wild type plants when grown under normal conditions (Fig. 5). This indicated that the EBE_TALE20_ mutation in the *MeSWEET10a* promoter of SC8 cultivar has no effect on yield-related traits in cassava.

**Fig. 5.**
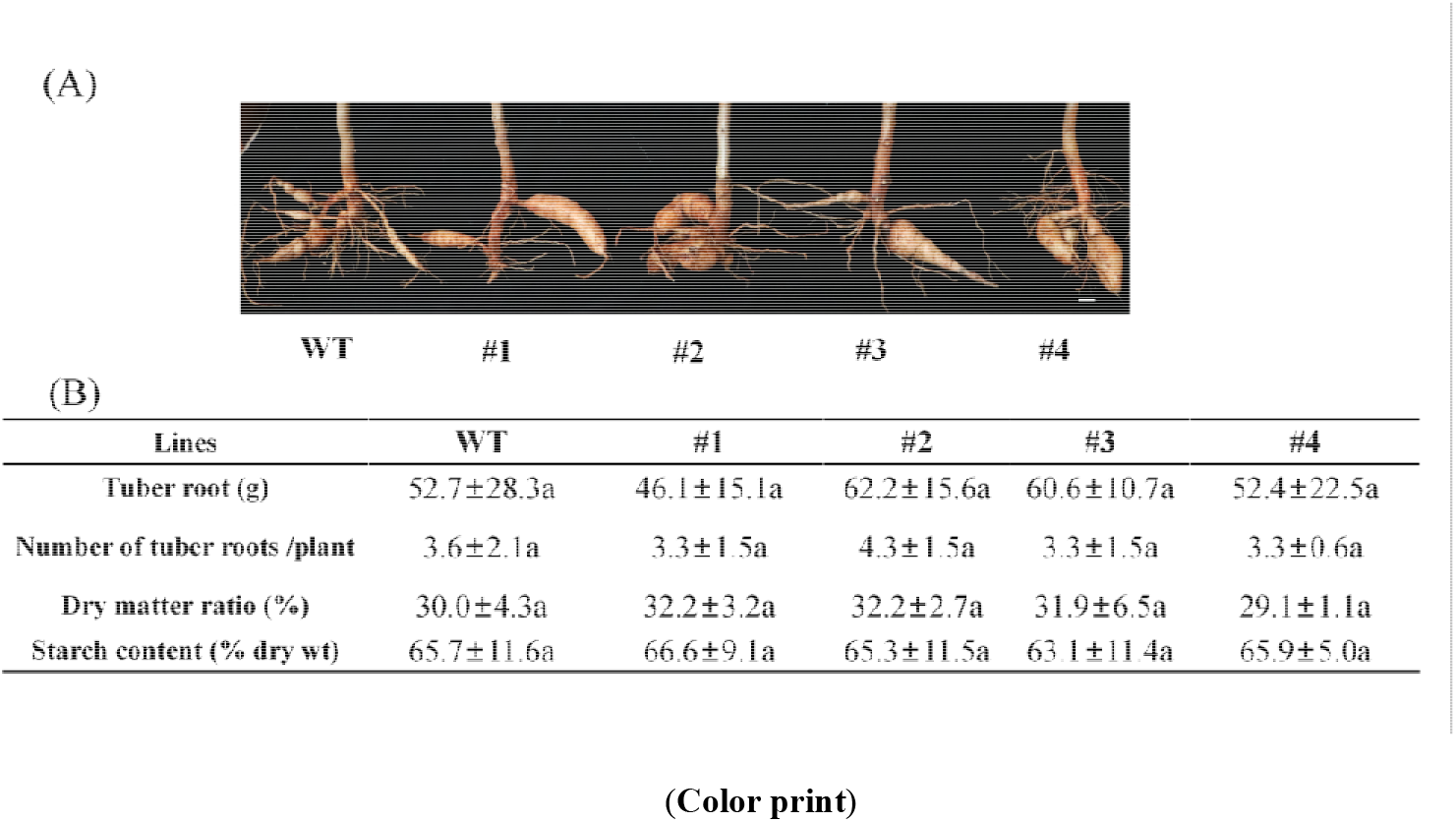
Tuber root phenotypic characterizations **(A)**. Scale bar, 1 cm. Yield-related traits shown as the mean ± *SD* from five plants of each line **(B)**. The values marked with the same letter (a) represented no significant difference (*p* < 0.01).

## 4. Conclusions

In conclusion, editing of the promoter in the disease-susceptibility gene *MeSWEET10a* in SC8 cassava confers resistance to CBB. All the mutated cassava lines had normal morphological and yield-related traits as the wild type. The results lay a research foundation for breeding Chinese local planting cassava SC8 cultivar resistant to bacterial blight by CRISPR/Cas9 technology.

## CRediT authorship contribution statement

**Yajie Wang**, **Mengting Geng** and **Ranran Pan:** Investigation, Conceptualization, Visualization, Writing – original draft. **Tong Zhang, Xiaohua Lu, Xinghou Zhen, Yannian Che, Ruimei Li** and **Jiao Liu:** Investigation, Validation. **Yinhua Chen**, **Jianchun Guo** and **Yao Yuan:** Writing - review & editing, Funding acquisition, Project administration, Supervision, Validation.

## Declaration of Competing Interest

The authors declare that they have no known competing financial interests or personal relationships that could have appeared to influence the work reported in this paper.

## Acknowledgments

This research was supported by the National Key Research and Development Program of China: 2019YFD1001105; the Hainan Provincial Natural Science Foundation of China: 320RC492; the Major Science and Technology plan of Hainan Province: ZDKJ2021012; Hainan Yazhou Bay Seed Lab: B21HJ0303 and the the earmarked fund for CARS: CARS-11.

## Notes

### Competing Interest Statement

The authors have declared no competing interest.

### Summary of Updates

Provides new data and experimental methods.

